# Complex history of aerobic respiration and phototrophy in the Chloroflexota class Anaerolineae revealed by high-quality draft genome of *Ca*. Roseilinea mizusawaensis AA3_104

**DOI:** 10.1101/2020.11.30.404129

**Authors:** Lewis M. Ward, Fátima Li-Hau, Takeshi Kakegawa, Shawn E. McGlynn

**Affiliations:** Department of Earth and Planetary Sciences, Harvard University, USA; Earth-Life Science Institute, Tokyo Institute of Technology, Japan; Department of Geosciences, Tohoku University, Japan

**Keywords:** Photosynthesis, thermophile, sulfide, phylogenetics, metagenomics

## Abstract

The candidate genus Roseilinea is a novel lineage in the Chloroflexota (formerly Chloroflexi or green nonsulfur bacteria) phylum known so far only from very incomplete metagenome-assembled genomes and preliminary enrichments. Roseilinea is notable for appearing capable of anoxygenic photoheterotrophy despite being only distantly related to well-known phototrophic Chloroflexota in the Chloroflexia class such as *Chloroflexus* and *Roseiflexus.* Here, we present a high quality metagenome-assembled genome of a member of Roseilinea, greatly improving our understanding of the metabolic capacity and taxonomic assignment of this genus. These data allow us to confidently describe the genetic basis for photoheterotrophy in these organisms as well as propose a candidate family for these organisms, *Ca.* Roseilineaceae, within the Anaerolineae class of Chloroflexota.

## Introduction

Anoxygenic phototrophs in the Chloroflexota (formerly Chloroflexi or Green Nonsulfur Bacteria) phylum include the well known genera *Chloroflexus* and *Roseiflexus* isolated and characterized from sulfidic hot springs (e.g. Thiel et al. 2018), but more recent metagenomic analyses have revealed several additional lineages of phototrophic Chloroflexota from environments including iron-rich hot springs (e.g. Ward et al. 2019, Ward et al. 2019b), carbonate tidal flats (Ward et al. 2020), and soda lakes (e.g. Grouzdev et al. 2018). While some of these novel phototrophic Chloroflexota belong to the Chloroflexia class together with *Chloroflexus* and *Roseiflexus* (e.g. Grouzdev et al. 2018, Ward et al. 2019c), several others belong to the poorly characterized Anaerolineae class (e.g. Klatt et al. 2011, Ward et al. 2019a). Most phototrophic Anaerolineae belong to several discrete lineages within the Aggregatilineales order (Ward et al. 2020); however, the first lineage of phototrophic Chloroflexota outside of the Chloroflexia class ever discovered—*Candidatus* Roseilinea gracile, first observed via metagenomic sequencing of a hot spring microbial mat from Yellowstone National Park (Klatt et al. 2011, Tank et al. 2017)—has never been confidently assigned a taxonomic rank below the class level. The relatively low quality of available genomes related to *Ca.* Roseilinea gracile, together with the absence of a pure culture isolate of this strain, has precluded thorough characterization and confident taxonomic assignment of this novel group of phototrophic Chloroflexota. Here, we present a high-quality draft genome of a bacterium within the *Ca.*

Roseilinea genus, dubbed here *Ca.* Roseilinea mizusawaensis AA3_104. This genome, recovered from metagenomic sequencing of Mizusawa Onsen in Japan, provides additional high-quality genomic data for characterizing the metabolic potential, evolutionary history, and taxonomic relationships of the Roseilinea genus. We propose here the assignment of Roseilinea to a novel family Roseilineaceae within the Chloroflexota order Thermoflexales, with *Ca.* Roseilinea mizusawaensis AA3_104 as type genome.

## Methods

Samples were collected from Mizusawa Hot Spring in Akita Prefecture, Japan. The sampling location was at 39°45’23.4”N 140°46’42.3”E where the sulfidic hot spring flowed down a steep gulley. The hot spring water was measured as pH 5.6 at the time of sampling. Samples consisted of microbial biomass collected as mottled green and white streamers in areas of flowing water at 50°C and a several mm thick, leathery microbial mat with white, green, red, and orange layers in 43°C water (Figure 1). Two samples from each site were selected for sequencing.

**Figure 1:**
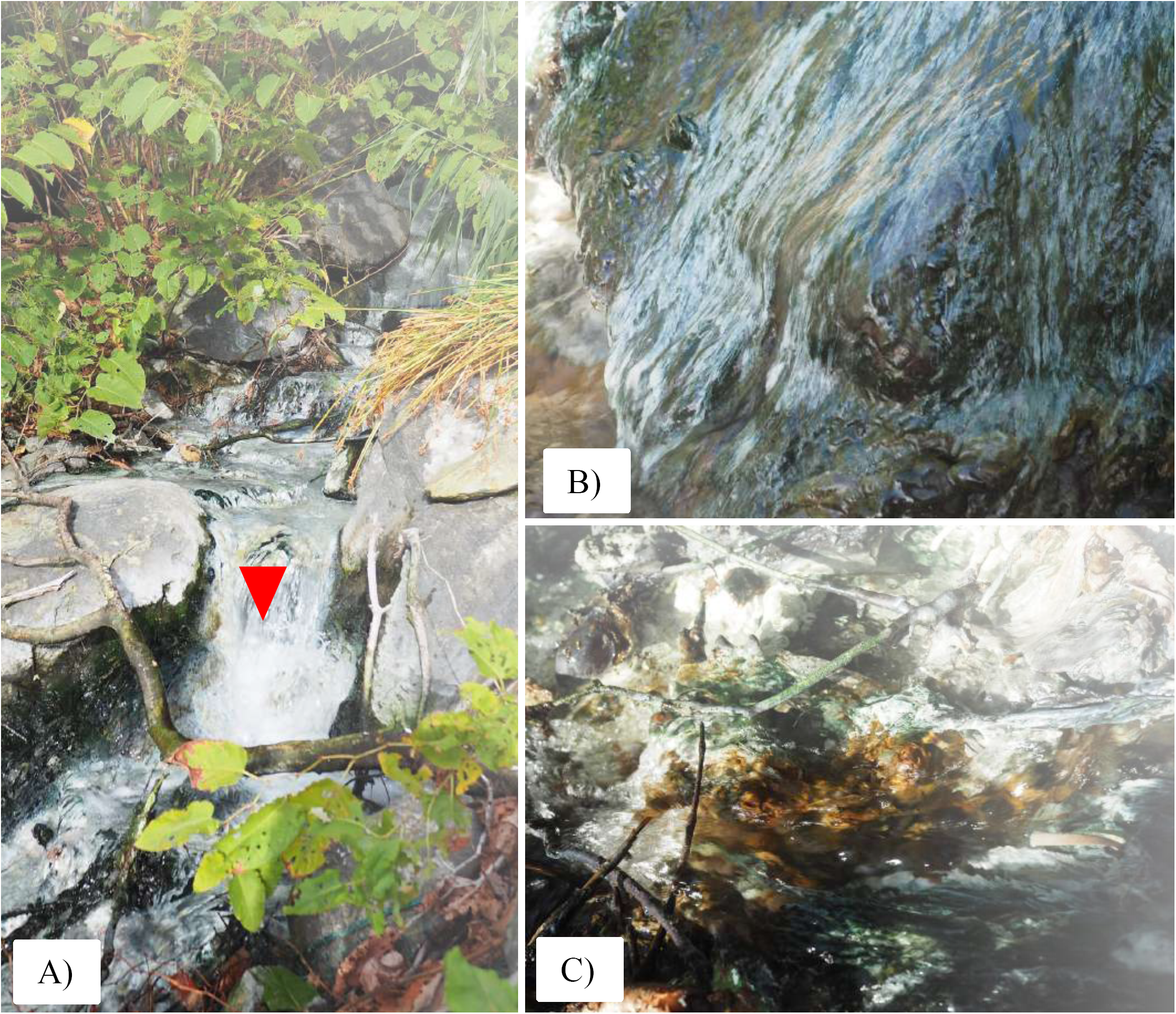
A) Overview of Mizusawa Hot Spring. B) White streamers in area of high flow. C) Thick laminated mat with multiple colored layers.

pH was measured with an Extech DO700 8-in-1 Portable Dissolved Oxygen Meter (FLIR Commercial Systems, USA). Temperature was measured with a Lasergrip 774 IR Thermometer (Etekcity, USA) and a ThermoPop thermometer (ThermoWorks, UK). The presence of sulfide was determined by smell.

Lysis of cells and preservation of DNA was accomplished in the field with Zymo Terralyzer BashingBead Matrix and Xpedition Lysis Buffer. Samples of microbial mat were disrupted immediately after sampling by attaching sample tubes to a cordless reciprocating saw (Makita JR101DZ) and operating for 1 minute. Bulk environmental DNA was extracted and purified after return to the lab with a Zymo Soil/Fecal DNA extraction kit. Quantification of DNA was performed with a Qubit 3.0 fluorimeter (Life Technologies, Carlsbad, CA) according to manufacturer’s instructions. Purified DNA was submitted to the Integrated Microbiome Resource for library preparation and sequencing following established protocols (Comeau and Langille 2017) with 2×150bp Illumina NextSeq. Raw sequence reads were quality controlled with BBTools (Bushnell 2014) and coassembled with MegaHit v. 1.02 (Li et al. 2016). Genome bins were constructed based on differential coverage using MetaBAT (Kang et al. 2015). Completeness and contamination/redundancy were determined with CheckM v1.1.2 (Parks et al. 2015). The genome was uploaded to RAST v2.0 for annotation and characterization (Aziz et al. 2008). Presence or absence of metabolic pathways of interest was predicted using MetaPOAP v1.0 (Ward et al. 2018b). Taxonomic assignment was determined with GTDB-Tk v1.2 (Parks et al. 2018, Chaumeil et al. 2019, Parks et al. 2020). Genomes were compared with AAI (Rodriguez and Konstantinidis 2014) to verify species and genus-level divisions. Organismal phylogenies were built using concatenated ribosomal proteins following methods from Hug et al. (2016). Protein sequences were extracted from genomes using the *tblastn* function of BLAST+ (Camacho et al. 2009) and aligned using MUSCLE (Edgar 2004). Trees were calculated using RAxML v8.2.12 (Stamatakis 2014) on the Cipres science gateway (Miller et al. 2010). Transfer bootstrap support values were calculated by BOOSTER (Lemoine et al. 2018), and trees were visualized with the Interactive Tree of Life viewer (Letunic and Bork 2016). All software was run with default parameters.

## Results and Discussion

Metagenomic sequencing of 4 samples from Mizusawa Onsen produced 119132162 reads totaling 5755889978 bp. Coassembly of these data produced 881712 contigs totaling 842365641 bp with and N50 of 1467 bp. AA3_104 was recovered as a metagenome-assembled genomes from binning of this dataset.

The AA3_104 MAG was recovered as 3855811 nt in 59 contigs with an N50 of 137185 nt. The genome is 61.6% GC and encodes 49 RNAs and 3396 coding sequences. The genome was estimated to be 97.71% complete, 1.1% redundant/contamination, and 0% strain heterogeneity by CheckM. The MAG encodes 49 RNAs and complete 16S, 23S, and 5S rRNA gene sequences. This MAG meets current standards for a high-quality draft genome (Bowers et al. 2017). Raw metagenomic reads mapped to the AA3_104 genome were recovered with ~1.7-fold coverage in the two samples from streamers collected from 50°C water and at up to 13.2-fold coverage in the samples from microbial mats from 43°C.

Phylogenetic analysis using concatenated ribosomal protein phylogenies (Figure 2) clustered AA3_104 with *Ca.* Roseilinea gracile and J036, enigmatic phototrophic Chloroflexota only distantly related to other phototrophs in this phylum. *Ca.* Roseilinea gracile and its relatives have previously been proposed to belong to the Anaerolineae (Klatt et al. 2011), *Ca.* Thermofonsia (Ward et al. 2018), and a novel order-level lineage (Ward et al. 2019). Taxonomic assignment of AA3_104, *Ca.* Roseilinea gracile, and J036 by GTDB-Tk determined that these organisms belongs to the Thermoflexales order within the Anaerolineae class of the Chloroflexota phylum, but could not place these genomes in a defined clade at the family rank or below. Together with their apparent phylogenetic divergence from other defined Chloroflexota orders suggests that these organisms represent a novel family within the Thermoflexales order of the Anaerolineae class of Chloroflexota. These genomes are 60-63% similar by AAI and 75-88% similar by ANI, consistent with these genomes representing three distinct species within a single genus (referred hereafter collectively as *Roseilinea*).

**Figure 2:**
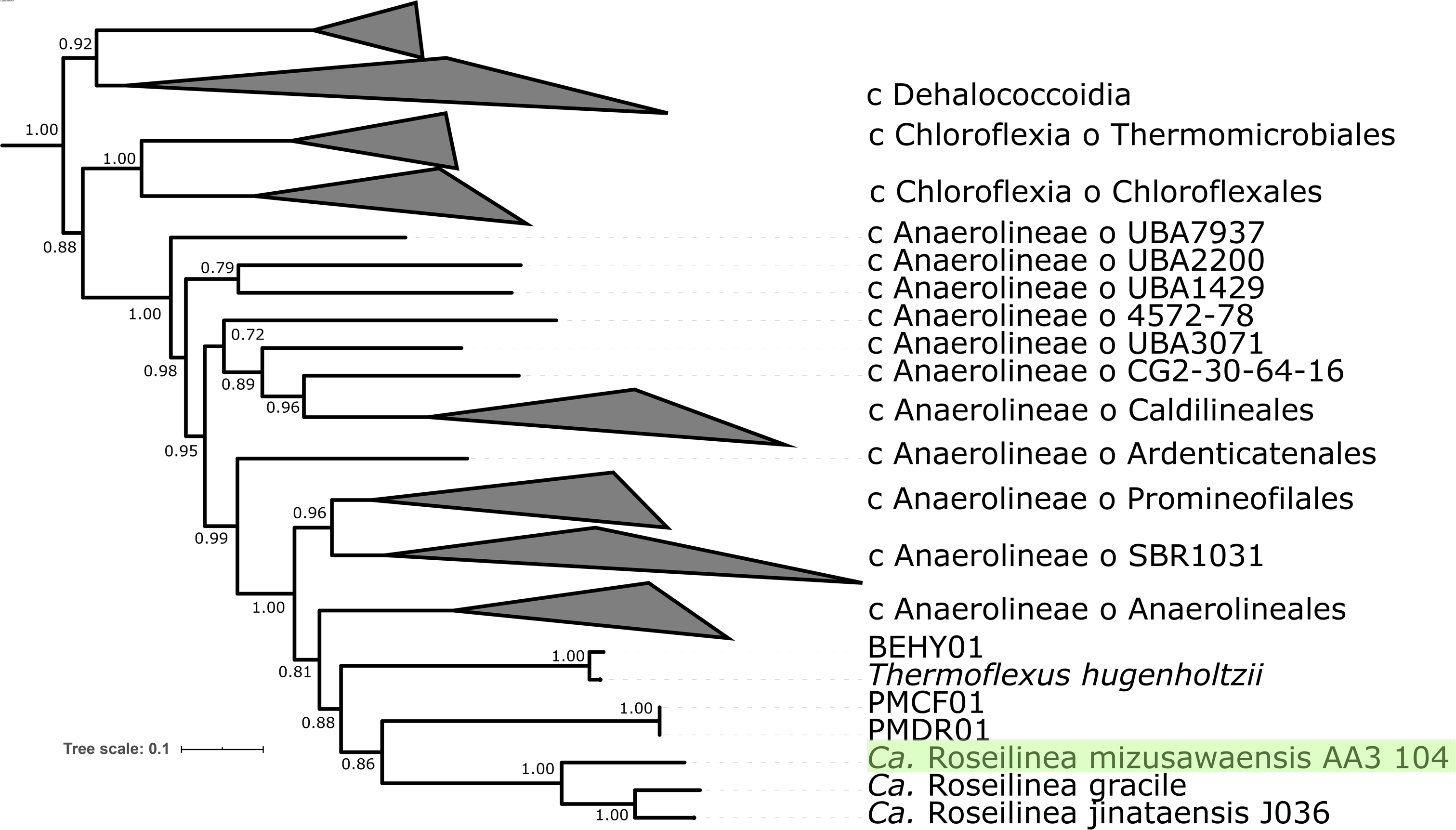
Phylogeny the Chloroflexota, built with concatenated ribosomal proteins following Hug et al. 2016 and rooted with the closely related phyla Eremiobacterota (Ward et al. 2019) and Armatimonadetes (Ward et al. 2017). The Thermoflexales order of the Anaerolineae class is shown at the strain level, while other clades are collapsed at the order (Chloroflexia and Anaerolineae classes) or class level.

Analysis of the metabolic potential of AA3_104 suggests similar capacities to other members of *Roseilinea;* importantly, however, the relatively high completeness and quality of the AA3_104 MAG relative to its relatives allows more confident determination of what proteins and pathways are or are not actually encoded by these organisms.

Like previously described *Roseilinea* strains, AA3_104 appears to be capable of photoheterotrophy via a Type 2 Reaction Center encoded by unfused *pufL* and *pufM* genes. AA3_104 appears to encode synthesis of bacteriochlorophyll *a* through an atypical pathway previously proposed for phototrophic Anaerolineae in which *bchM, bcheE,* and the *bchLNB* complex are absent (e.g. Ward et al. 2018). The high completeness of AA3_104 (>97%) greatly increases confidence that the apparent absence of these genes is not a false negative caused by failure to recover sequences in the MAG (MetaPOAP false negative probability <10^-9^). Known alternatives to missing bacteriochlorophyll synthesis genes (e.g. *acsF* for *bchE,* genes encoding the light-dependent POR enzyme for genes encoding the BchLNB complex, Chew and Bryant 2007) were also absent. Despite the apparent absence of several components of the classical bacteriochlorophyll synthesis pathway from *Roseilinea* genomes, observations of cells putatively identified as *Ca.* Roseilinea gracile do show autofluorescence consistent with bacteriochlorophyll a (Tank et al. 2017), suggesting that this pigment is in fact synthesized in these organisms. Genes such as *bchK, bchU,* and *bchQ* are absent from AA3_104, consistent with this organism not producing bacteriochlorophylls c, d, or e. Taken all together, these data are consistent with previous proposals that phototrophic Anaerolineae (both *Roseilinea* and members of the order Aggregatilineales) utilize a novel bacteriochlorophyll a synthesis pathway that makes use of uncharacterized or multifunctional enzymes to perform steps typically performed by BchE, BchM and the BchLNB complex (e.g. a multifunctional BchXYZ complex for the conversion of both protochlorophyllide a to chlorophyllide a and the conversion of chlorophyllide a to 3-vinyl-bacteriochlorophyllide a) (Ward et al., 2018; Ward and Shih 2020).

Like previously described phototrophic Anaerolineae (including *Roseilinea* and phototrophic members of Aggregatilineales), but unlike phototrophic Chloroflexia, AA3_104 lacks key marker genes for carbon fixation pathways including the Calvin cycle (e.g. rubisco) and the 3-hydroxypropionate bi-cycle (e.g. the trifunctional malyl-CoA/beta-methylmalyl-CoA/citramalyl-CoA lyase). AA3_104 is therefore most likely incapable of autotrophy and instead makes a living as an anoxygenic photoheterotroph.

AA3_104 appears to be capable of at least facultative aerobic respiration via both an A- and a B-family heme copper O_2_ reductase (HCO), two *bc* Complex IIIs, and a *bd* oxidase. This complement of respiration genes is consistent with that previously described for members of *Roseilinea* (Ward et al. 2018, Ward et al. 2019).

The presence of a B-family HCO in phototrophic *Roseilinea* is a trait shared with all other known phototrophic Chloroflexota as well as phototrophic members of the closely related phylum Eremiobacterota, although closely related nonphototrophic strains lack a B-family HCO (e.g. Ward et al. 2018, Ward et al. 2019b, Ward et al. 2020). The functional link between the B-family HCO and phototrophy in these organisms is not well understood, but may relate to the adaptation of B-family HCOs to relatively low oxygen concentrations (Han et al. 2011) and the oxygen sensitivity of proteins involved in anoxygenic phototrophy (e.g. Hamilton 2019) tying anoxygenic photoheterotrophy in these organisms to low-oxygen environments. Interestingly, phylogenetic relationships of B-family HCO proteins are incongruent with organismal relationships and with relationships among phototrophy proteins (Figure 3, Figure 4). This suggests that B-family HCOs and phototrophy proteins have independent histories of horizontal gene transfer in the Chloroflexota despite their apparent functional link.

**Figure 3:**
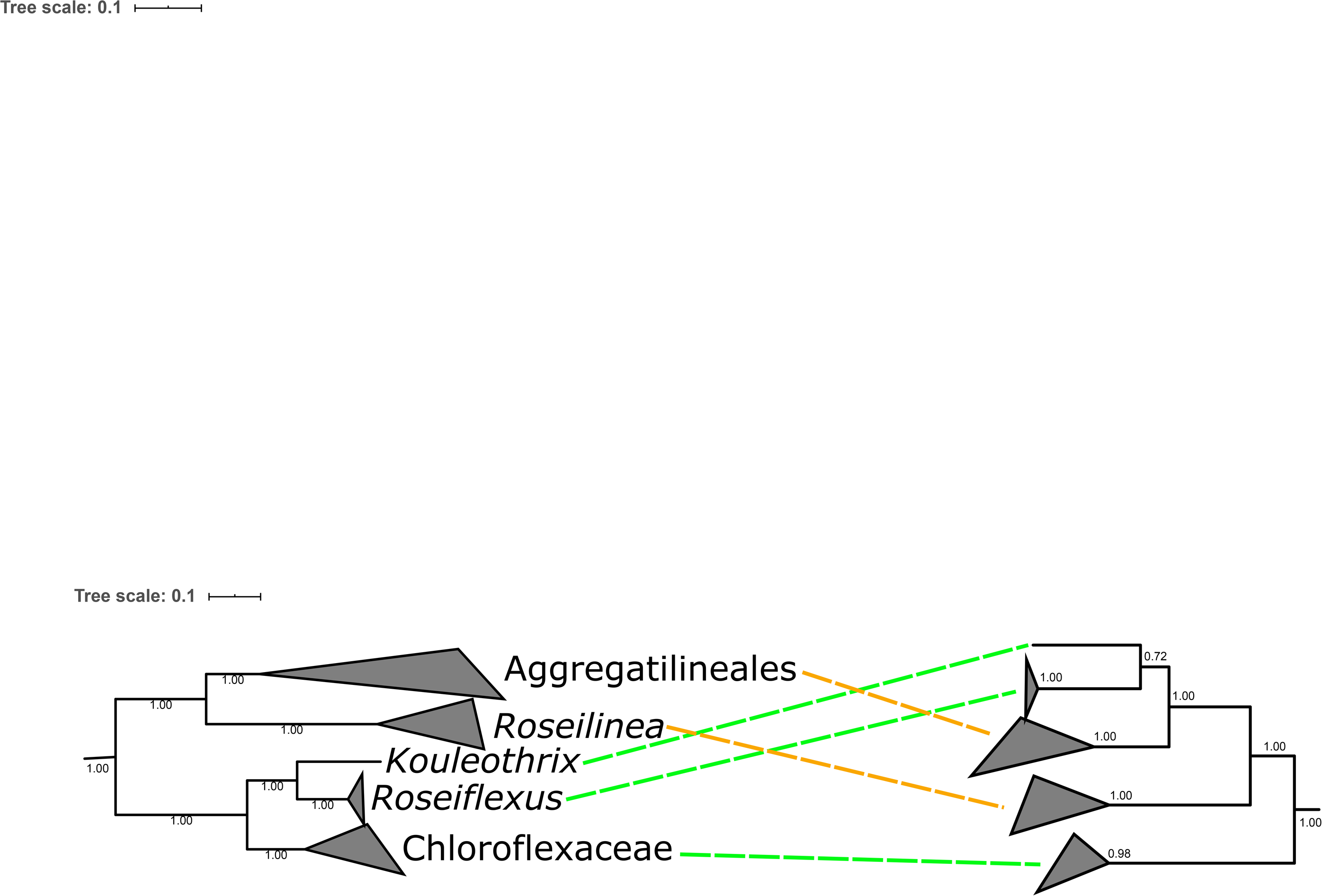
Tanglegram demonstrating phylogenetic incongruity between organismal phylogeny (left, built with concatenated ribosomal proteins) and phototrophy protein phylogeny (right, built with concatenated PufL, PufM, BchX, BchY, and BchZ sequences) of phototrophic Chloroflexota, collapsed at the genus (*Roseilinea, Roseiflexus,* and *Kouleothrix*), family (Chloroflexaceae) or order (Aggregatilineales) levels. While phototrophy proteins appear to have been vertically inherited within the Chloroflexia class after the divergence of the basal nonphototrophic Herpetosiphonaceae family (Ward et al. 2015, Ward et al, in press), the incongruent positioning of phototrophic Aggregatilineales and *Roseilinea* suggests that these lineages acquired phototrophy via horizontal gene transfer from members of the Chloroflexia (Ward et al. 2018). Secondary HGT events may have been occurred within Aggregatilineales (Ward et al. 2020)

Unlike other phototrophic Chloroflexi, AA3_104 and other *Roseilinea* do not encode an Alternative Complex III. While the absence of genes encoding this protein from previous MAGs could have been ascribed to their low completeness, the high quality of AA3_104 makes it quite unlikely that this enzyme would not have been recovered in the MAG if it were found in the source genome (MetaPOAP false negative estimate <2.5% for AA3_104 alone, <10^-4^ for all three available *Roseilinea* genomes considered together). It therefore appears that members of *Roseilinea*, in contrast to other phototrophic Chloroflexota, use a *bc* Complex III in their phototrophic electron transport chain instead of an Alternative Complex III. The evolutionary and biochemical logic for this difference is not yet apparent, but it does appear to confirm that an Alternative Complex III is not essential for phototrophy in the Chloroflexota.

In most lineages of phototrophic bacteria, it appears that the capacity for aerobic respiration was acquired before the acquisition of phototrophy (Fischer et al. 2016). This trend has been confirmed for phototrophs in the Chloroflexota orders Chloroflexales and Aggregatilineales, in which A-family HCOs and other components of electron transport chains were acquired to enable aerobic respiration before the acquisition of phototrophy and B-family HCOs (Shih et al. 2017, Ward et al. 2020). However, at present it is impossible to confirm whether this trend extends to *Roseilinea.* While genes for aerobic respiration are widespread in even apparently obligately anaerobic members of the Anaerolineae class of Chloroflexota (e.g. Hemp et al. 2015, Pace et al. 2015, Ward et al. 2018b), it appears that these genes were acquired independently in many lineages within this class subsequent to their divergence (e.g. Ward et al. 2020). Members of Roseilineaceae described so far consist of a single apparent genus at the end of a relatively long branch whose closest relatives appear to be a family of Thermoflexales provisionally identified by GTDB as Fen-1058 (Figure 2). Fen-1058 does encode aerobic respiration; however, respiratory proteins in the families of Thermoflexaceae—Roseilineaceae, Fen-1058, and Thermoflexaceae—are not closely related (Figure 4). This suggests that each of the known families of Thermoflexales acquired respiration independently after they diverged. Both aerobic respiration and phototrophy therefore appear to have been acquired along the long branch leading to crown group *Roseilinea;* in the absence of additional information it is therefore impossible to determine which of these traits was acquired first along this branch. Resolving this uncertainty will require recovering additional Roseilineaceae diversity that breaks up the long branch leading to *Roseilinea.* However, it is clear from phylogenetic relationships of proteins that components of the electron transport chain in Roseilineaceae were acquired through independent HGT events from different sources, suggesting that the capacity for respiration and phototrophy was not acquired in a single large HGT event.

**Figure 4:**
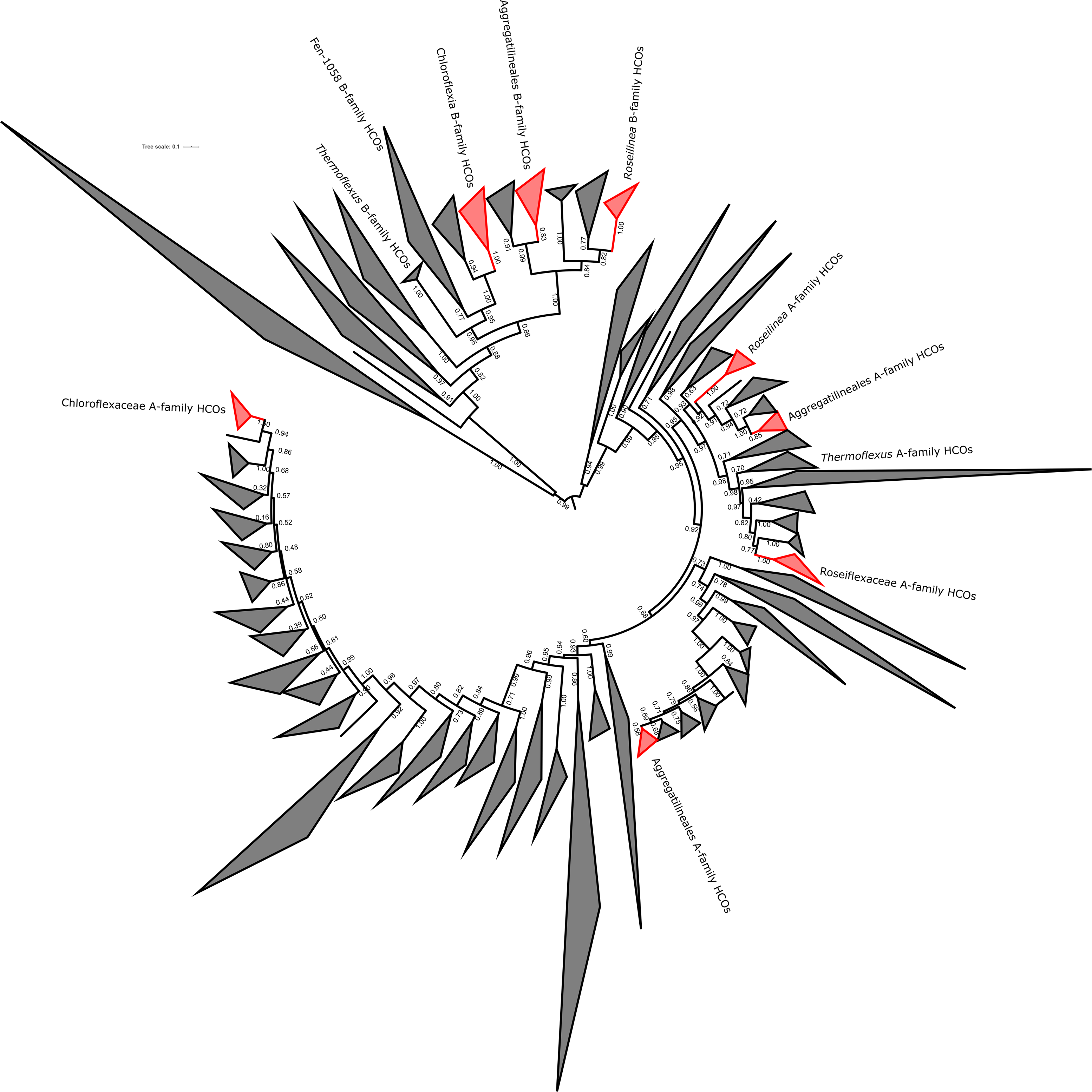
Phylogeny of Heme-Copper Oxidoreductase (HCO) protein sequences with clades consisting of sequences from phototrophic Chloroflexota highlighted in red and labeled. While all known phototrophic Chloroflexota encode both an A- and a B-family HCO for O2 reduction, these appear to have been acquired independently, with HCO phylogenies incongruent with both organismal relationships as well as with relationships between phototrophy proteins. Clades containing HCOs from other members of Thermoflexales are also labeled to highlight their phylogenetic distance from those of *Roseilinea.*

## Conclusions

We propose the assignment of AA3_104, described here, and J036, described in Ward et al. 2019, to the *Roseilinea* genus proposed by Tank et al. (2017). We propose the specific epithets *Ca.* Roseilinea mizusawaensis for AA3_104 in recognition of Mizusawa Onsen as the source of this organism, and *Ca.* Roseilinea jinataensis for J036 in recognition of its discovery in Jinata Onsen. Given the apparent divergence of *Roseilinea* from other members of the phylum Chloroflexota as determined by concatenated ribosomal protein phylogenies as well as analysis via GTDB-Tk, we propose the assignment of these organisms to a novel family in the Thermoflexales order of the Anaerolineae class of the Chloroflexota phylum, Roseilineaceae, fam. nov.. As it is currently the highest quality MAG available for this clade, we propose *Ca.* Roseilinea mizusawaensis AA3_104 as the type genome for Roseilineaceae until such time as an isolate and/or a complete genome is available.

The family Roseilineaceae is so far known only as putatively at least facultatively aerobic photoheterotrophs from hot springs. Members of Roseilineaceae are currently known from geographically and geochemically diverse hot springs in the United States and Japan, including iron-rich and moderately acidic intertidal hot springs in Tokyo Prefecture (Ward et al. 2019), moderately acidic and sulfidic hot springs in Akita Prefecture (this study), and alkaline siliceous hot springs in Yellowstone National Park (Klatt et al. 2011). However, 16S rRNA gene amplicon data suggests that this clade may have a wider environmental distribution. Highly similar 16S sequences (>97%) to that of *Ca.* Roseilinea mizusawaensis AA3_104 have been reported from wastewater treatment systems (Karst et al. 2018) while somewhat similar (>94%) sequences have been reported from environments including contaminated aquifers (Thavamani et al. 2012) and soil (Xie et al. 2009). Given the long phylogenetic branch between crown group *Roseilinea* and the divergence of Roseilineaceae from *Thermoflexus* and other members of Thermoflexales, together with current understanding of the diversification rates of bacterial lineages through time (Louca et al. 2018), it should be expected that much more diversity of Roseilineaceae has existed through geologic time. Ignoring possibilities of apparently rare extreme extinction events and population bottlenecks, it seems highly likely that additional Roseilineaceae lineages exist in the environment today. Additional genome-resolved metagenomic sequencing of diverse environments is likely to yield additional diversity of this group, potentially breaking up the relatively long phylogenetic branch leading to *Roseilinea* and allowing for better resolution of the evolutionary transitions that have led to the significant divergence of *Roseilinea* from its closest known relatives.

*Ca.* Roseilinea mizusawaensis AA3_104 improves the available genomic diversity of anoxygenic phototrophic Chloroflexota outside of the Chloroflexia class, and provides the best available metagenome-assembled genome within the newly proposed Roseilineaceae family (Table 1). Improving sampling across the diversity of Chloroflexota—particularly within novel phototrophic lineages—provides valuable insight into the evolutionary trends leading to the extant diversity of phototrophs within this phylum and across the entire tree of life. In particular, the apparently extensive role of horizontal gene transfer in shaping the distribution of phototrophy and related pathways across the tree of life (e.g. Raymond et al. 2002, Hohmann-Marriott and Blankenship 2011, Fischer et al. 2016, Ward et al. 2018a, Shih et al. 2017, Ward et al. 2019b, Ward and Shih 2019) provides an opportunity to investigate the history and evolution of functional traits even though most bacteria that have possessed these traits through geologic time are now extinct (Louca et al. 2018).

**Table:**
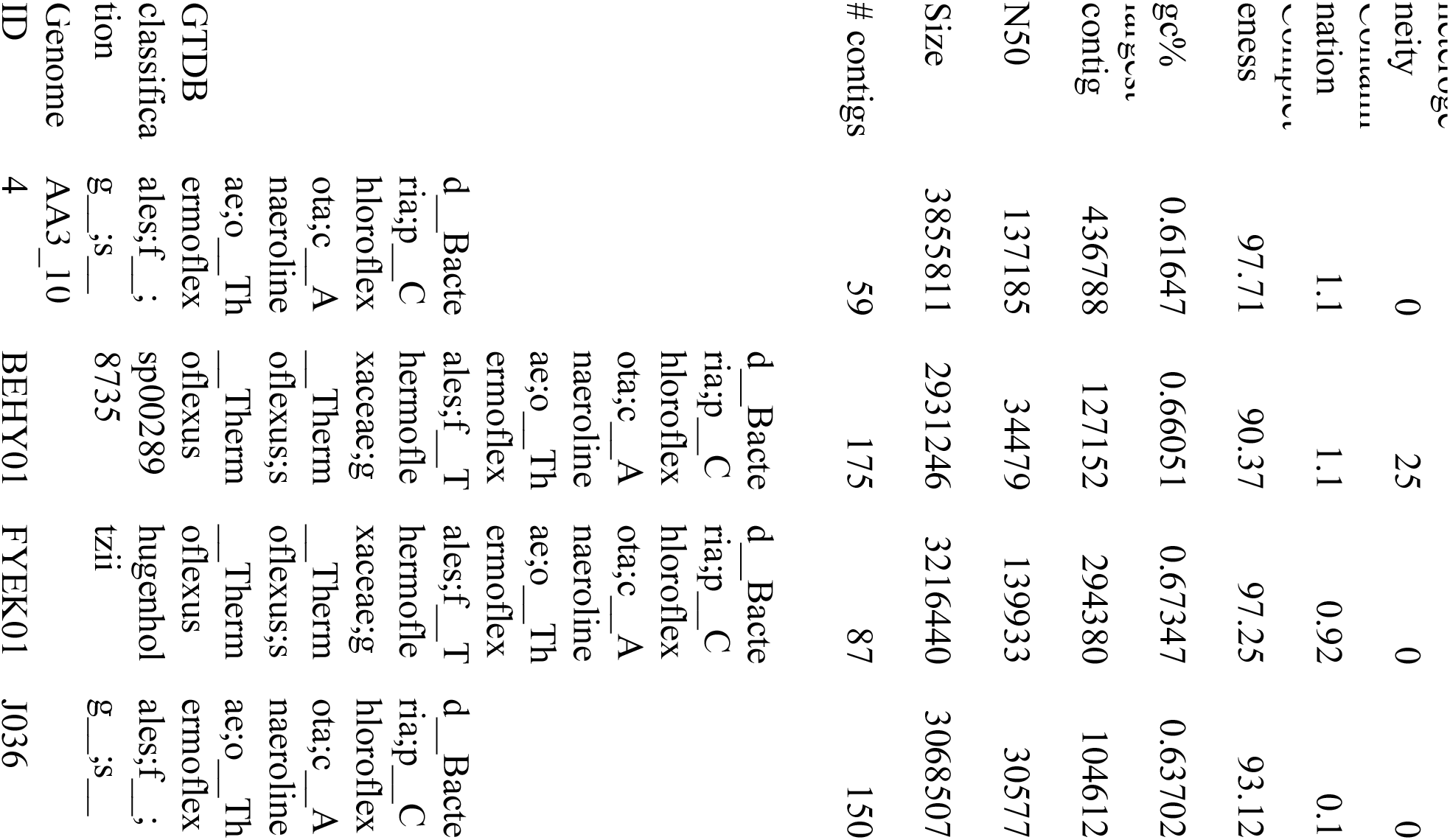
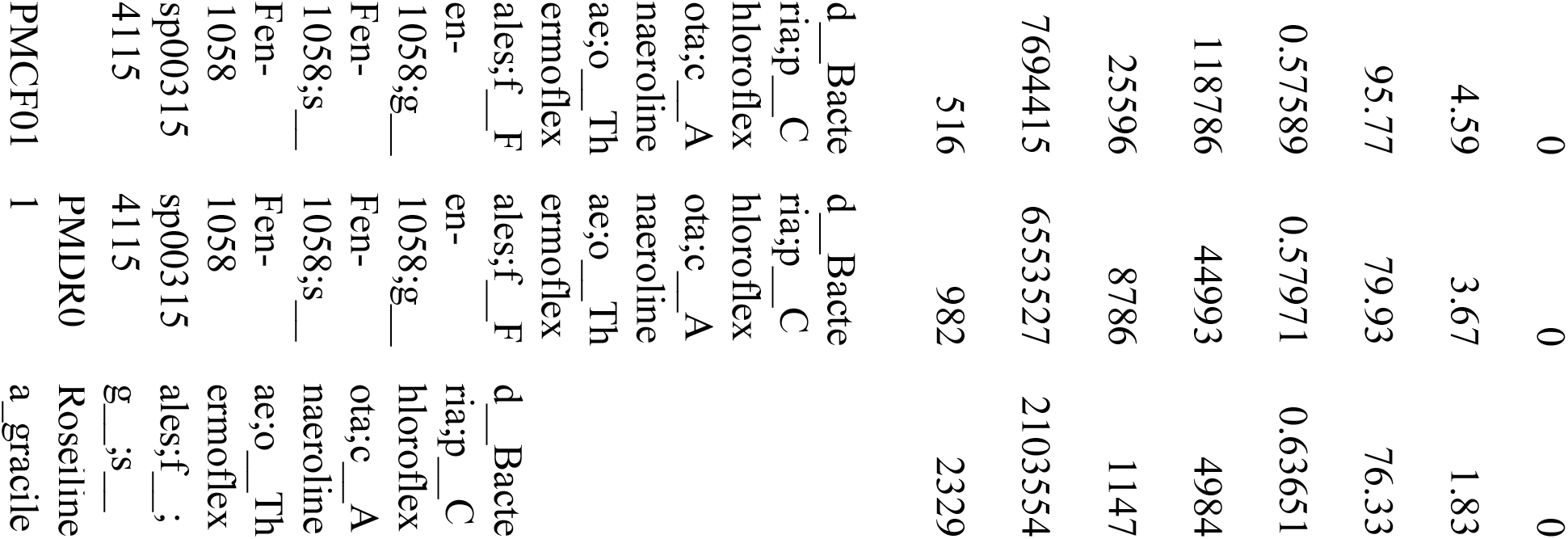
Genome statistics and GTDB-Tk taxonomic assignments of *Roseilinea* and other Thermoflexales strains.

Utilizing the extant diversity and evolutionary relationships of microorganisms to understand the early evolution of metabolic traits requires adequate sampling across extant diversity. The recovery of *Ca.* Roseilinea mizusawaensis AA3_104 and other members of Roseilineaceae provide an excellent example of the taxonomic and metabolic novelty that exists in the environment which can be discovered via genome-resolved metagenomic sequencing and which will provide crucial data for comparative phylogenetic analyses that can help answer longstanding questions about the evolution of phototrophy, respiration, and other traits deep in the tree of life.

## Data Availability

The AA3_104 genome has been uploaded to the NCBI WGS database under the submission ID SUB8655613 and will be publicly available immediately following processing.

## Author Contribution Statement

All authors performed fieldwork and read and approved the manuscript. LMW and FLH processed samples. LMW performed analyses and prepared the manuscript. LMW and FLH prepared figures.

## Acknowledgements

LMW acknowledges support from a Simons Foundation Postdoctoral Fellowship in Marine Microbial Ecology. SEM acknowledges support from the Astrobiology Center Program of the National Institutes of Natural Sciences (grant no. AB311013).

